# Understanding and exploitation of host stress responses to protein production using novel transcriptomic analytics

**DOI:** 10.1101/2023.11.30.569415

**Authors:** Christoffer Rode, Felix Beulig, Mathias Jönsson, Leonie J. Jahn, Morten H. H. Nørholm, Bernhard O. Palsson, Emre Özdemir, Lei Yang

**Affiliations:** Novo Nordisk Foundation Center for Biosustainability, Technical University of Denmark, Kemitorvet, Building 220, 2800, Kgs. Lyngby, Denmark; Department of Bioengineering, University of California, San Diego, La Jolla, USA

**Keywords:** Big data, Transcriptomics, Protein production, iModulon analysis, Strain design, Stress response

## Abstract

Predictable expression of heterologous genes in a production host is a fundamental challenge in biotechnology. While traditional methods focus on manipulating expression and the property of the heterologous gene, a systems biology approach can complement with designs to improve the host itself. Previously, Independent Component Analysis (ICA) of the RNAseq data helped reveal independently modulated gene sets (iModulons) in bacteria. This was later applied to identify common stress responses related to heterologous gene expression for *Escherichia coli*. In this study, we expand this analysis with additional non-enzymatic proteins and apply our findings to design novel protein production optimization. By leveraging the Precise-1K transcriptomics knowledge base, we identify three iModulons as novel transcriptional responses to protein production stress; Cold Shock, gcvB sRNA, and the uncharacterized UC-9 iModulons. By studying the gene membership in the UC-9 iModulon, we discover effective novel design targets for improving protein production. This study demonstrates the value of big data analytics and systems understanding of host responses for designing novel strategies to optimize protein production.

## Highlights

● Big transcriptomic data analytics reveals stress responses in production hosts.
● Stress from heterologous genes depends on the gene itself and its expression level.
● Cold Shock, gcvB, RpoH, and Rsp iModulons are activated host responses.
● An uncharacterized iModulon is now associated with protein production stress.
● Co-overexpression of *rspAB* and choline supplementation can increase protein production.

## 1. Introduction

Microbial hosts are widely used for recombinant protein production in biotechnology for manufacturing of both high-value pharmaceutical proteins (Puetz and Wurm, 2019; Rettenbacher et al., 2022) and low-cost industrial enzymes (Arnau et al., 2020; Kirk et al., 2002). Other industrial proteins currently emerging include food ingredient proteins, sweet-tasting proteins, structural proteins, and antifreeze proteins, etc. (Augustin et al., 2023; Eskandari et al., 2020; Miserez et al., 2023). Despite various strategies available to improve product titers and folding (Davy et al., 2017), optimizing protein production in microbial hosts remains a challenge.

Many studies have investigated how -omics data can be leveraged to identify optimization strategies (Chen et al., 2020). Current -omics technologies have advanced greatly in recent years and can provide systems-level knowledge about production hosts. For example, RNA sequencing (RNAseq) has been used to understand how *E. coli*, a popular production host (Pouresmaeil and Azizi-Dargahlou, 2023), copes with protein production stress at the transcriptional level. For certain proteins, the host upregulates heat shock stress functions and downregulates energy metabolism and carbon source utilization genes (Dürrschmid et al., 2008; Haddadin and Harcum, 2005; Oh and Liao, 2000), and may upregulate osmoprotectant uptake and mRNA degradation genes (Sharma et al., 2011). This has been used to guide strain design; by co-overexpressing heat shock stress genes, host viability and protein production can be improved (Ow et al., 2010). However, these studies have been limited to specific proteins, and a systematic overview of host responses to different proteins is still lacking. Big data analytics offers tools to obtain a deep understanding of complex - omics data. For large transcriptomics datasets, this has been accomplished through independent component analysis (ICA). Here, gene expression data is decomposed into independently modulated sets of genes known as iModulons (Sastry et al., 2019). A knowledge base has been developed for transcriptomics datasets for different organisms (iModulonDB), and recently, a large library of 1035 high-quality *E. coli* samples was assembled, known as Precise-1K (Lamoureux et al., 2023; Rychel et al., 2021). Before this, ICA was used on *E. coli* transcriptomes to study transcriptional responses to overexpression of 40 heterologous enzymes (Tan et al., 2020). Here, iModulons were identified that commonly changed activities among all the samples, including fear vs. greed, metal homeostasis, and protein folding responses.

In this study, we further expand the understanding of responses related to protein production by overexpressing an additional set of 10 genes encoding for non-catalytic proteins. By applying the large Precise-1K RNAseq compendium, we find that the transcriptomes of strains overexpressing different heterologous genes can be clustered according to their transcriptional profiles. Each cluster shares some major changes to specific iModulons, including novel responses not revealed in previous observations (Tan et al., 2020). One such hallmark iModulon is the previously uncharacterized iModulon UC-9. We further investigated the genes in this iModulon and successfully implemented design strategies to improve protein production. This work provides a workflow for translating systems-level knowledge into practical strain or process design.

## 2. Methods and Materials

### 2.1 Strain Construction

The *Escherichia coli* MG1655 variant Gly2 was selected for this study. This strain grows optimally on glycerol and contains the following mutations: *yegE* S683Y*, glpK* L65M*, rpoC* L770R (Tan et al., 2020). We further modified this host with a knockout of the *rhaBADM* operon to prevent consumption of rhamnose that was used to induce expression from plasmids. The knockout was introduced by CRISPR/MAD7-mediated Lambda Red recombination. Primers and linear DNA fragments used for cloning are provided in Supplemental Data 1. pGE3 containing the MAD7 endonuclease and the Lambda Red Exo, Beta, and Gamma genes were transformed into a Gly2 strain and selected on ampicillin. Then, pSD128 containing a gRNA targeting rhaD and a crRNA compatible with the MAD7 endonuclease was used to introduce a double-stranded break in the *rhaBADM* operon and was transformed into the cells together with the SD_PR443 repair oligo. Cells were selected on ampicillin and chloramphenicol. The final strain was cured from plasmids and counterselected on minimal M9 media with rhamnose as the sole carbon source to verify the knockout. Its genome was sequenced revealing the Δ4,500 bp deletion of the *rhaBADM* operon and an additional Δ16,400 bp spontaneous deletion of a transposable element between genes *yhiM* – *yhiS*.

### 2.2 Assembly of plasmid library

A library of expression plasmids was generated for the overexpression of heterologous genes using primers and linear DNA fragments provided in Supplemental Data 1. For the list of heterologous gene sequences, see Supplemental Data 2. The expression plasmid is based on a pNIC28-Bsa4 backbone modified with a *PrhaSR-PrhaBAD* inducible promoter system. The backbone was amplified from pSD51 used by Tan et al. to express the MBP protein (Tan et al., 2020). All heterologous genes were codon-optimized (Shen and Packer, 2022), and inserted downstream of an identical BCD sequence and a His-TEV tag via gibson assembly. Heterologous genes were ordered as gBlocks from Integrated DNA Technologies with overhangs complementary to the cloning site of the vector backbone. An empty plasmid was used for control without the cloning site and *PrhaBAD* promoter sequence and was cloned via USER cloning. Heterologous gene expression was also tested in a T7-based expression system, where the T7 polymerase gene was expressed from a bacterial artificial chromosome and the heterologous gene was inserted into the multiple cloning site in a pET-52B vector by USER cloning. For the assembly of *rspAB* co-overexpression plasmids, an expression site consisting of a promoter (BBa_J23117), the *rspAB* operon, and a terminator (t500) was introduced next to the heterologous gene expression site. This was done by amplifying vector backbones with primers modified with overhangs to promoter and terminator sequence. The *rspAB* operon was amplified from MG1655 genomic DNA using primers modified with complementary overhangs. Vector and insert were subsequently assembled via USER cloning.

### 2.3 Shake flask cultivations

All cultivations were carried out under the same growth conditions. Cells were grown in 250 mL shake flasks at 37°C under 225 RPM in 40 mL minimal M9 medium. Glycerol (0.2% v/v) was used as the sole carbon source and media was supplemented with appropriate antibiotics, Wolfe’s vitamin solution (0.1% v/v), and an iron-rich trace metal solution (0.05% v/v), see Supplemental Table S1 for its composition. Main cultures were inoculated from a 1:100 dilution of pre-cultures (15 mL in 250 mL shake flasks) that had been growing overnight between 16-17 hours. These pre-cultures were inoculated from a 1:100 dilution of minimal M9 media glycerol stocks of OD = 1. The main cultures were induced at OD 0.40 with rhamnose (final conc. 1 mM). Strains with a T7-based expression system was induced with IPTG (final conc. 1 mM). Main cultures were sampled for RNAseq and western blot analysis 2 hours post-induction.

### 2.4 Western blotting

Protein production was verified by Western Blots (Figure S1). Cell pellets were lysed by heating (95°C for 15 minutes) in a reducing buffer consisting of 90% Invitrogen Laemmli buffer (Bio-Rad) and 10% NuPAGE (Invitrogen). Lysate was loaded on precast Bio-Rad Protean SDS-PAGE gels for 5 minutes at 120 V, then 30 minutes at 200 V. After gel electrophoresis, proteins were transferred onto an iBlot membrane using an iBlot 2 Western Blot Transfer System (Thermo Fisher). Blots were incubated in skim milk solution (50 g/L skim milk powder in TBS-T) for 1 hour at 4°C, then washed for 5 minutes in TBS-T over three rounds. Primary his tag antibody (05-949, Merck), stored in a skim milk solution, was added and incubated overnight at room temperature. The primary antibody was then removed and replaced with a secondary HRP antibody (GENA9310, Cytiva) in skim milk solution and incubated for 1 hour at room temperature. Chemiluminescence was then developed on the blots by adding ECL Western Blotting Detection reagents (Cytiva) and incubating for 3 minutes in darkness.

### 2.5 Flow cytometry

Fluorescence was measured by flow cytometry by first diluting cell cultures in a PBS buffer and then injected on an Agilent NovoCyte Quanteon Flow Cytometer System. To filter out debris, threshold settings used for forward scatter and side scatter thresholds were set to 7.500 and 5.000, respectively. No gates were used. Fluorescence intensity levels were then calculated as the mean value of FITC-A measurements for 10,000 events.

### 2.6 Transcriptomics

An RNAseq library was generated in biological duplicates. After 2 hours of induction, RNA was extracted from cultures following the protocol in the RNAprotect Bacteria Reagent Handbook (Qiagen). A culture volume equal to 3 mL of OD = 1 was added to 6 mL RNAprotect Bacteria Reagent (Qiagen), vortexed, and incubated at room temperature for 5 minutes. Sample pellets were resuspended in 400 µL elution buffer and split into two aliquots which were stored at -80°C. Total RNA was then extracted and purified from thawed pellets on a QIAcube Connect Instrument using the RNeasy Protect Bacteria Mini Kit (Qiagen).

Lysozyme was used for enzymatic lysis of cells together with Proteinase K for degrading protein and SUPERase•In RNAse inhibitor (Thermo Fisher) for protecting RNA. An optional DNase treatment step was followed, and cells were eluted in 30 µL RNase-free water, which was stored at -80C. Thawed total RNA was depleted from rRNA using the Ribo-Zero rRNA Removal Kit for Gram-Negative Bacteria (Illumina) according to the protocol. RNA libraries were prepared for paired-end sequencing using the KAPA Stranded RNA-Seq Library Preparation kit (Kapa Biosystems). Libraries were sequenced on a NextSeq system (Illumina) using the QIAseq FastSelect -5S/16S/23S kit (Qiagen). The transcriptomic reads were mapped to the combined sequence of the host genome and expression plasmid sequence by following the Modulome workflow (https://github.com/avsastry/modulome-workflow, commit: f15878d). Here, reads were normalized to transcripts per million (TPM), and then log-transformed for RNA-seq analysis. Metadata of samples and log-TPM data can be found in Supplemental Data 3 and 4. The relative levels of heterologous mRNA (Figure 1b) were calculated as heterologous gene counts over the total gene counts.

**Figure 1.**
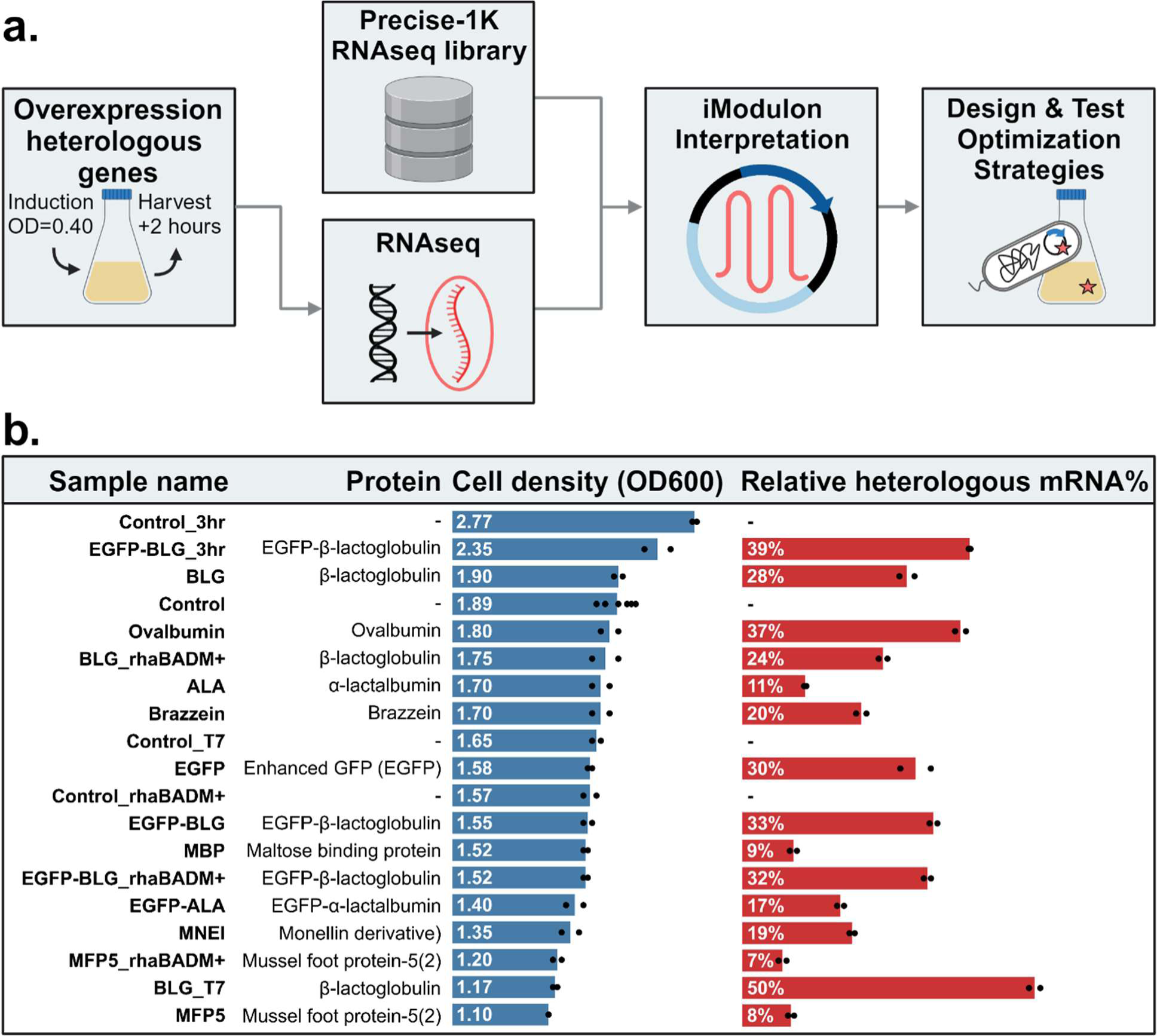
Workflow and library of samples used to understand and address transcriptional stress responses. **a.** Workflow for generating and analyzing transcriptomics data. **b.** Description of transcriptome samples including sample names, proteins, cell density (OD600) at sampling time, and the relative percentage of heterologous mRNA, calculated as the heterologous gene counts over the total gene counts.

### 2.7 Combining transcriptomics data with Precise-1K transcriptomic datasets and inferring iModulon activities

Log-TPM data was batch corrected using pyComBat (Behdenna et al., 2023), then inspected for quality and appended to the Precise-1K RNAseq compendium detailed in https://github.com/SBRG/Precise1k-analyze (Lamoureux et al., 2023). This resulted in a total of 1,081 expression profiles, 46 of which were generated for this study. Genes on expression plasmids were removed for this analysis and native gene expression data was centered on the Precise-1K wildtype controls (samples: control wt_glc 1 and control wt_glc 2). The iModulon activities in our new dataset were inferred from the Precise-1K using the M matrix. For the analysis, iModulons related to genetic differences were filtered out. This was done by removing iModulons under the system categories Genetic alterations (22 iModulons) and ALE effects (13 iModulons) based on iModulon annotations from Precise-1K (Lamoureux et al., 2023).

### 2.8 Revising the UC-9 iModulon by independent component analysis

We further characterized the UC-9 iModulon for potential gene memberships and gene weights by performing an Independent Component Analysis (ICA) on the combined dataset of PRECISE-1K and our newly generated RNAseq dataset. The FastICA algorithm was implemented using Scikit-learn (v0.20.3) with 100 iterations and a convergence tolerance of 1e-7 (Sastry et al., 2019). To determine the ideal number of components, we used the OptICA method, with a maximum of 370 dimensions and a step size of 20 (McConn et al., 2021). We repeated the entire process 100 times to ensure that the final calculated components were robust. The resulting iModulons and the gene membership in the revised UC-9 iModulon, Rsp, were characterized through the Pymodulon workflow (https://github.com/SBRG/pymodulon/) (Sastry et al., 2019).

## 3. Results and Discussion

### 3.1 Workflow for studying heterologous gene overexpression stress at the transcriptional level in *Escherichia coli*

To understand how overexpression of heterologous genes stresses production hosts, we generated a transcriptomics dataset for the overexpression of 10 different heterologous genes encoding non-enzymatic proteins that vary in size and structure listed in Supplemental Data 2. These include bovine whey proteins β-lactoglobulin (BLG) and α-lactalbumin (ALA), chicken egg white protein ovalbumin, two sweet-tasting proteins brazzein (Ming and Hellekant, 1994) and monellin derivative MNEI (Leone et al., 2016), a dimerized Mussel Foot Protein-5 (MFP5) (Danner et al., 2012; Kim et al., 2018), the native well-folding Maltose Binding Protein (MBP) (Tan et al., 2020), and an enhanced green fluorescent protein (EGFP). We also designed and included fusion proteins, EGFP-BLG and EGFP-ALA, with EGFP fused at the N-terminal, intended to screen for optimized protein production hosts. These heterologous genes were expressed from an expression system using the native RNA polymerase (RNAP). For comparison, a strong T7-RNAP expression system was tested for BLG (BLG_T7). A workflow was designed where RNAseq samples were taken in exponential phase during heterologous gene overexpression in *E. coli* (Figure 1a), described in Methods and Materials 2.3. A total of 45 RNAseq samples were included for twenty different gene overexpression or control conditions, done in biological duplicates (Figure 1b).

The controls in this study were benchmarked with many other available datasets by comparing iModulon activities with the wildtype controls in Precise-1K (see Figure S2). This analysis showed a list of iModulons that are differentially activated in our controls. RpoS and ppGpp iModulons have lowered activity in this study, which may result from the strain and growth settings we defined. Most of the other differential iModulons can be attributed to the composition of the medium. The Glycerol iModulon is activated because glycerol was used as the carbon source. Iron uptake also showed low activity in this study due to a higher than standard concentration of iron in our trace metal solution (see Methods and Materials). Metal regulation iModulons were generally amongst the most differentially activated iModulons in our controls. To obtain a better reference of our defined growth conditions, we also tested adjustments to certain samples. Controls sampled one hour later (Control_3hr) showed higher RpoS iModulon activity (stringent response), see Figure 2a. In contrast, Controls without the knockout of the rhamnose degradation pathway (Control_rhaBADM+) showed lower RpoS and an active Rhamnose iModulon. The control containing the T7 gene (Control_T7) showed no large changes in iModulon activities.

**Figure 2.**
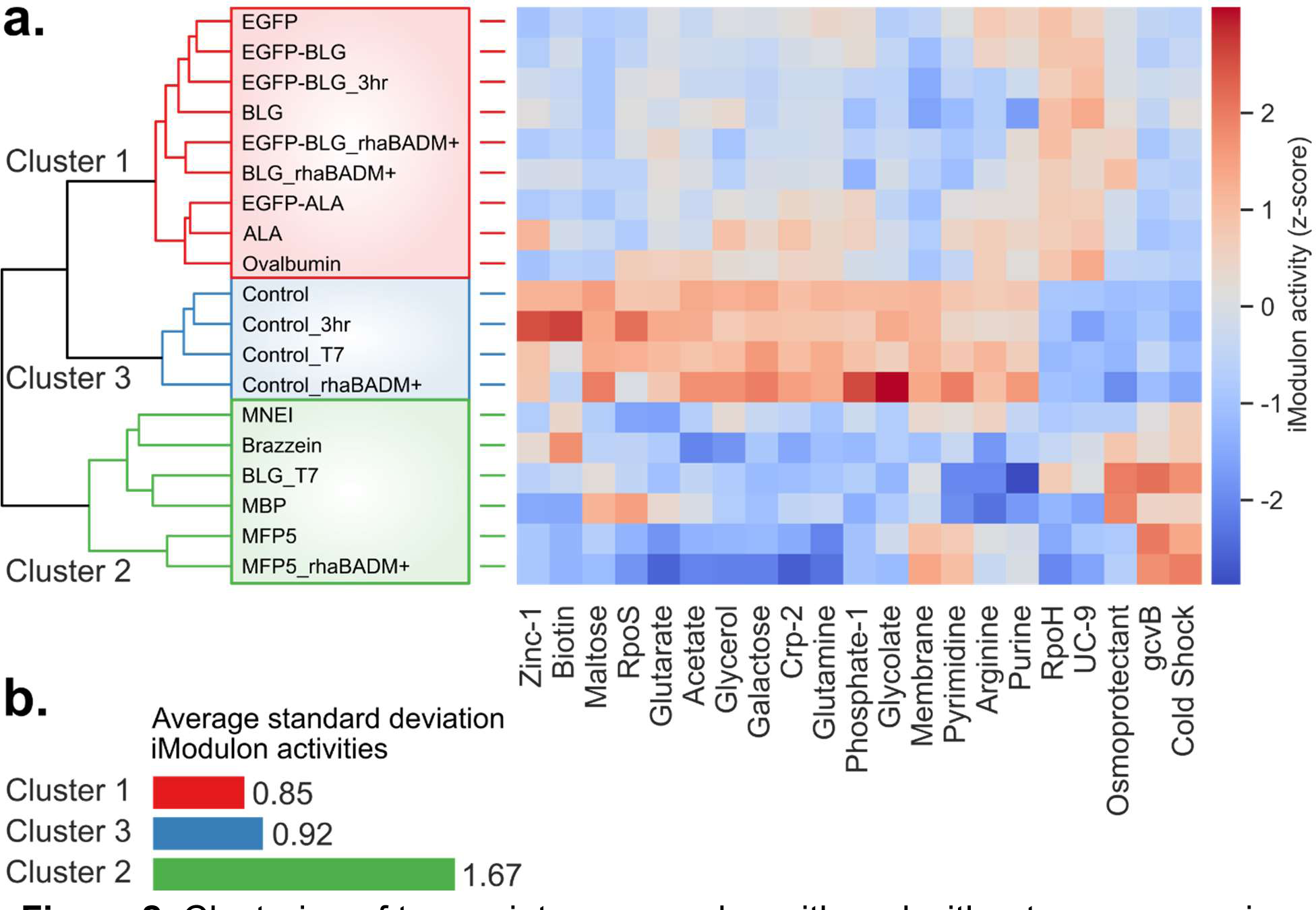
Clustering of transcriptome samples with and without overexpression of heterologous genes. **a.** Clustering of all samples based on gene expression data. The top 21 differentially activated iModulons during overexpression conditions and their standardized activities are shown in the heatmap (right). **b.** Average standard deviation for all iModulon activities within Cluster 1, 2 and 3.

Heterologous gene expression generally decreased growth, as shown in Figure 1b. Unexpectedly, strains expressing BLG and ovalbumin genes reached similar cell densities as the control. However, the expression of the larger EGFP fusion gene caused a drop in final densities. Production of MFP5, a tyrosine-rich protein with a repetitive sequence resulted in the lowest biomass in our samples, suggesting that characteristics of the target protein affect growth (Figure 1b). Strains with intact rhamnose metabolism genes, *rhaBADM*, did not reach higher cell densities. We found that relative heterologous mRNA levels differed markedly depending on the overexpressed gene, despite the use of the same promoter, bicistronic design and N-terminal tag. Using the strong T7-based expression system yielded higher heterologous mRNA levels (50% of total mRNA reads), compared to the rhamnose system (28%), but with the trade-off of 61% less final biomass (final OD600 1.90 against OD600 1.18). Results indicate that the T7-based expression exerts a substantial burden on the host.

### 3.2 The mode of host response depends on the overexpressed gene

To obtain an overview of transcriptional profiles in the various production strains, we clustered the complete set of gene expression data hierarchically using Ward’s method, see Figure 2a. This produced three separate clusters. Cluster 1 and Cluster 2 (red and green respectively) contain all overexpression samples, while Cluster 3 (blue) contain all control samples, where no gene is overexpressed. Cluster 1 contains samples for production strains expressing genes encoding for BLG, EGFP-BLG, ALA, EGFP-ALA, ovalbumin, and EGFP. Cluster 2 contains the remaining samples for genes encoding for MNEI, MFP5, MBP, and Brazzein. The high T7-based gene expression sample of BLG (BLG_T7) would also be grouped into Cluster 2. Interestingly, EGFP fusion proteins showed similar transcriptional profiles to their non-fused partners. For example, the profile for ALA is similar to that of EGFP-ALA, despite EGFP being a larger protein than ALA.

We then investigated the set of iModulons that are most differentially activated during overexpression of the heterologous genes. This was calculated by taking the iModulon activities of overexpression samples (Cluster 1 and 2) and subtracting them from the activities of the control samples (Cluster 3). For the analysis of iModulons, we filtered out the Rhamnose iModulon as well as iModulons under the system categories Genetic Alterations (22 iModulons) and ALE effects (13), based annotations from Precise-1K (Lamoureux et al., 2023). The resulting top 21 differentially activated iModulons are shown in Figure 2a. We calculated the standard deviation of iModulon activities within the three clusters, shown in Figure 2b and expanded in Figure S3. As shown, samples in Cluster 2 show a high variation in iModulon activities, as compared to Cluster 1. This indicates that samples in Cluster 1 activate a similar host response. In general, all genes overexpressed in Cluster 1 samples are globular proteins, while the genes in Cluster 2 encode more diverse proteins. This includes fibrous proteins (MFP5), globular proteins that are controlled by a very strong promoter (BLG_T7), and proteins that are small (MNEI) and cysteine-rich (Brazzein). We speculate that if more proteins are included, we would see Cluster 2 be further broken into sub-clusters. From the data we collected (Supplemental Data 4) we were unable to find a clear correlation between protein features and the clustering, but a much larger library of diverse proteins may enable us to predict the transcriptional responses from the parameters of the target protein.

### 3.3 Overexpression of heterologous genes reduce stringent response and metabolism functions in the host

The most notable iModulons had lower activities compared to controls, see Figure 2a. Among these, the most downregulated iModulons are mainly related to Stringent response (RpoS), carbon metabolism (Glycerol, Maltose, Galactose, and Crp-2), Nucleotide metabolism (Pyrimidine and Purine), and Amino acid metabolism (Arginine and Glutamine). One consequence of heterologous gene expression is that such genes occupy a portion of the total mRNA pool upon induction, which decreases the relative levels of native gene transcripts. We found this effect to be substantial; for example, the heterologous gene in the BLG_T7 condition comprises 50% of the total mRNA pool (Figure 1b). Modulating the pool of native transcripts can allow the host to spend more of its translation machinery on heterologous mRNA, which has been exploited in other studies to increase protein production in hosts with reduced genomes (Lieder et al., 2015; Ziegler et al., 2021). Effective genome reduction could be achieved by tailoring the reduction design to align with the transcriptome that is being allocated in the production host. The RpoS iModulon, comprising a large set of 122 genes to provide the stringent response, is deactivated throughout all overexpression samples. This agrees with the previously observed inverse correlation between gene expression levels and RpoS iModulon activity, which is in contrast to intuition (Tan et al., 2020). Instead, resources were presumably allocated to other cellular functions to support overexpression of the heterologous genes. It has been shown that reducing RpoS protein levels or even knocking out the *rpoS* gene, can be beneficial for increasing protein production titers in *E. coli* during fed-batch fermentations (Jeong et al., 2004; Weikert et al., 2000).

Other transcriptional responses are differentially activated between overexpression samples in Groups 1 and 2. This is shown in Figure 3 by comparing iModulon activities with controls. Samples in Cluster 1 commonly activate two iModulons; the heat shock iModulon RpoH and the uncharacterized iModulon UC-9 (Figure 3a). Samples in Cluster 2 have diverse transcriptional profiles, but generally share activation of the Cold Shock and gcvB iModulons (Figure 3b). Among these findings, activation of UC-9, Cold Shock, and gcvB iModulons are novel compared to what has been reported by Tan et al. (Tan et al., 2020). We will further discuss these iModulons in the following sections.

**Figure 3.**
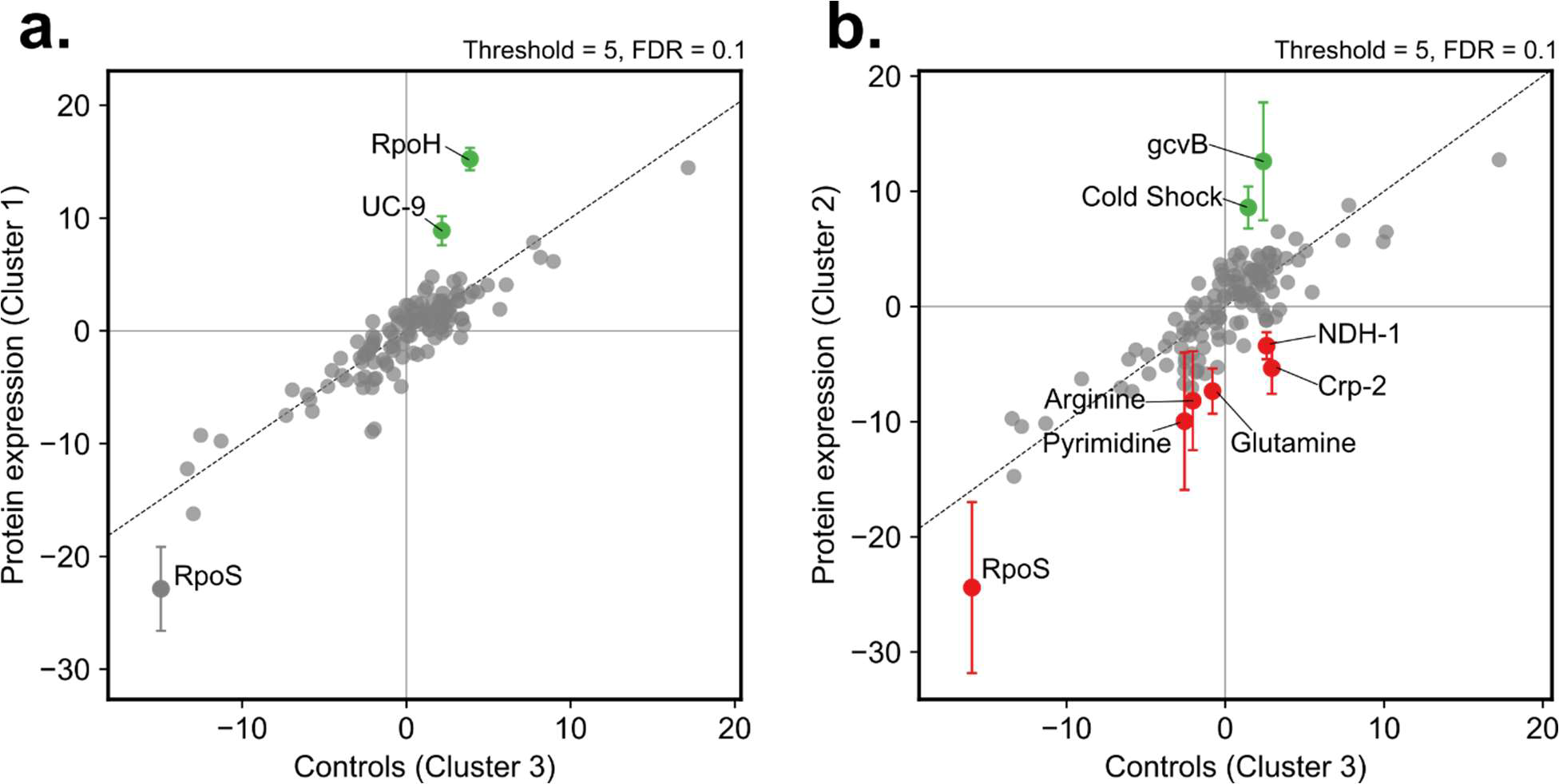
Differential iModulon Activity in overexpression samples of Cluster 1 (**a.**) and Cluster 2 (**b.**), compared to controls in Cluster 3. Significantly activated (green) or deactivated (red) iModulons are determined using Threshold = 10 and False Discovery Rate = 0.10 settings.

### 3.4 Activation of Cold shock and gcvB iModulons is common for overexpression samples in Cluster 2

Despite the large variation within Cluster 2 overexpression samples, there is a shared activation of the Cold Shock and gcvB iModulons. The function of gcvB iModulon is annotated in iModulonDB as related to amino acid transport. It has a main regulator gcvB, which is a small RNA critical for amino acid metabolism and transport (Lalaouna et al., 2019). However, the gcvB iModulon (76 genes) and gcvB regulon (79 genes) share only about 18% of genes (14 genes, all non-annotated), and about 70% of the genes in the gcvB iModulon are of unknown function (53 genes). Recently, it was also discovered that GcvB, together with RNA chaperone Hfq, regulates RNase BN by protecting its mRNA degradation from RNase E (Chen et al., 2019). This suggests that *gcvB* and related genes may have other functions outside of amino acid metabolism. For further investigation, the genes of unknown function in the gcvB iModulon need to be studied to understand this transcriptional response and its role in overexpression stress.

The Cold Shock iModulon includes cold shock proteins whose primary functions are to prevent the stabilization of unfavorable secondary DNA or RNA structures resulting from low-temperature conditions (Giuliodori, 2016; Jones and Inouye, 1994). The observation that Cold shock iModulon is activated during overexpression of heterologous genes is intriguing. It is generally beneficial to produce proteins at low temperatures in different microorganisms (Bhatwa et al., 2021; Rong et al., 2023). Besides temperature downshift, cold shock proteins can also be induced by other, less studied stress conditions. For example, chloramphenicol, an antibiotic that binds to ribosomes and inhibits translation, has been shown to induce the transcription of *cspA*, a major cold shock gene in *E. coli* (Jiang et al., 1993). At low temperatures, mRNA can form undesirable secondary structures that slow down the movement of ribosomes on mRNA and translation. CspA homologs can act as RNA chaperones to maintain single-stranded mRNA and facilitate translation (Phadtare, 2004). Cold shock proteins might be able to promote translation in a similar manner, even when cold shock genes are upregulated independently of lower temperature conditions. To test this, we focused on a specific protein, BLG. When the BLG encoding gene was expressed from the rhamnose-induced promoter, the strain was grouped in Cluster 1. However, when the gene was expressed from the T7 promoter, the transcriptomic profile had higher similarity with Cluster 2 overexpression samples. Expression from the T7 promoter resulted in a much higher fraction of heterologous mRNA and also a growth defect. Comparing the iModulon activities between these expression systems, surprisingly only 5 iModulons changed significantly (Figure S4b), including Cold shock and gcvB iModulons. We speculate that the strong T7 promoter outpaced the translation, thus leading to the induction of Cold shock iModulon, similar to CspA activation in response to chloramphenicol that inhibits translation. We further estimated the translation efficiency of the two constructs using a ribosome binding site calculator (Reis and Salis, 2020). The result showed that the translation initiation rate of the 5’UTR T7-based expression system is only a thirtieth the rate of the rhamnose expression system for the expression of BLG (Figure S4c), indicating a potential imbalance of transcription and translation for the T7-based expression system. The exact mechanism for Cold shock iModulon activation will need further investigation. Understanding the function of both Cold shock and gcvB iModulons is vital to identify effective strain design strategies that mitigate stresses from heterologous gene overexpression.

Despite these commonalities, the overexpression of the genes for Cluster 2 resulted in different and highly individualized transcriptomes. Details on differential iModulon activities for these transcriptomes are shown in Figure S5. Overexpression of the gene encoding the small and cysteine-rich Brazzein was found to activate a distinct set of iModulons that are related to redox stress and metal homeostasis – a response unlike any other of our tested heterologous gene. This agrees with previous reports that disulfide bond formation yields oxidative stress (Tyo et al., 2012). It is known that cytoplasm is not suitable for producing proteins requiring disulfide bonds, due to its reduced environment (Bhatwa et al., 2021). We speculate that cells producing Brazzein were not able to maintain the redox balance, leading to reduced growth (Figure 1). A more in-depth investigation of all iModulons categorized under Redox Stress, Cofactor Metabolism, and Metal Homeostasis is illustrated in Figure S6.

### 3.5 Overexpression samples in Cluster 1 activate the RpoH iModulon and the uncharacterized UC-9 iModulon

Overexpression of genes in Cluster 1 encoding for the globular proteins is distinguished by activation of the heat shock iModulon RpoH and an uncharacterized iModulon UC-9. About 83% of the genes in the RpoH iModulon (41 genes) are found in the RpoH regulon (140 genes), including heat shock proteins, chaperones, and proteases. These genes are regulated by the RpoH sigma factor (σ^32^) and are upregulated when protein misfolds or aggregates (Arsène et al., 2000).

Since the uncharacterized UC-9 iModulon is particularly active during overexpression samples in Cluster 1 (see Figure 3a), we investigated this iModulon further. The iModulon consists of 10 genes, but its activity is mainly driven by the two genes in the *rspAB* operon (Figure 4a). Intriguingly, co-overexpressing them has been shown to increase activity levels of overexpressed β-galactosidase (Weikert et al., 2000). However, the molecular function of RspA and RspB is ambiguous. One study showed that *rspAB* overexpression decreases RpoS levels (Huisman and Kolter, 1994), while a more recent study characterized RspA as a D-altronate dehydratase (Wichelecki et al., 2014).

**Figure 4.**
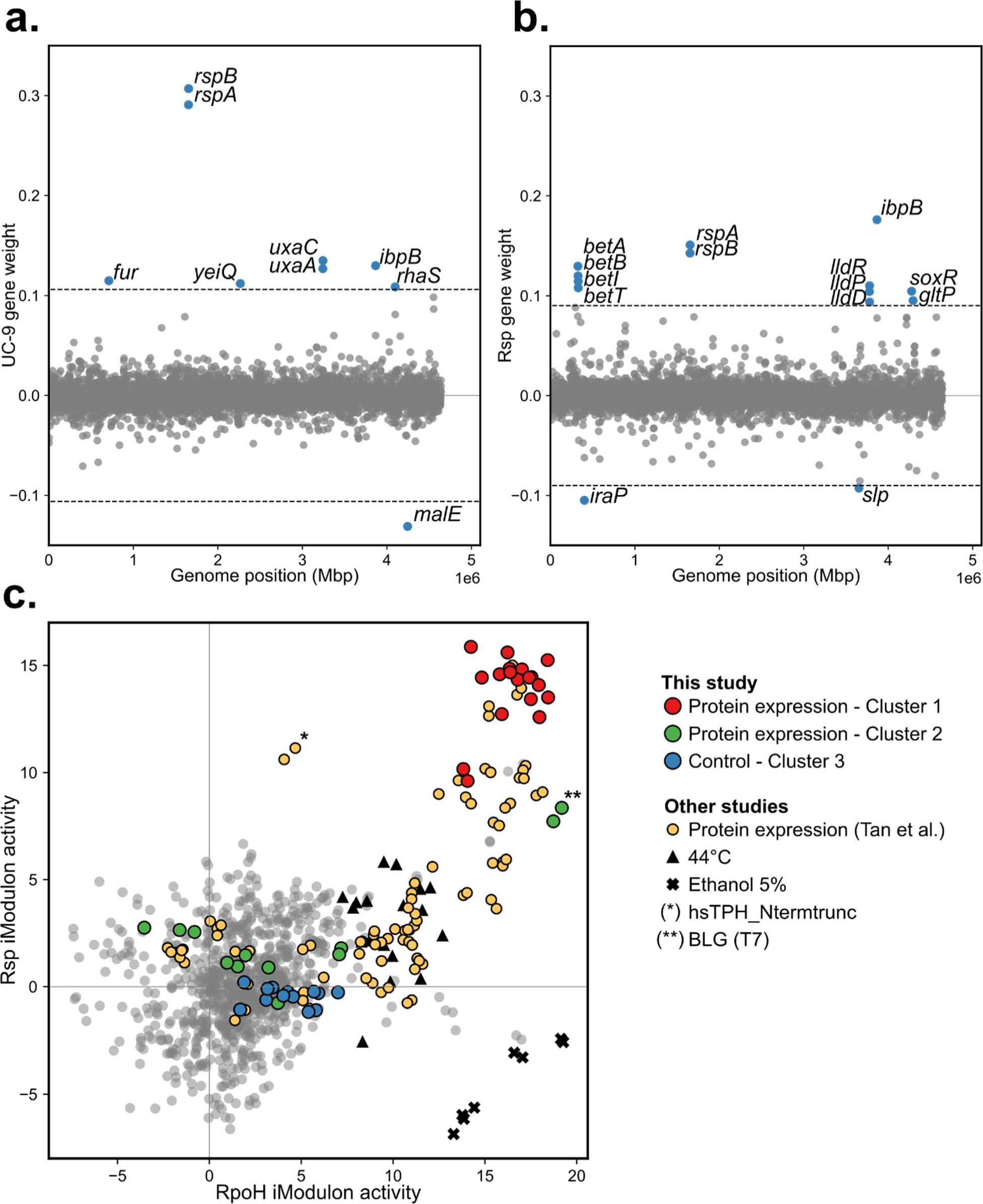
Gene membership of the Rsp iModulon and its positive correlation with the RpoH iModulon in Precise-1K samples. **a.** Gene membership of the UC-9 iModulon from Precise-1K. Only genes beyond the threshold contribute with weight to the iModulon activity. **b.** Gene membership in the revised Rsp iModulon was found by including our dataset with Precise-1K in the ICA. For clarity, the threshold was increased by 15% to remove low-weight genes from the figure. **c.** The link between the Rsp and RpoH iModulons for all samples in the Precise-1K RNAseq library.

To better understand the function of the UC-9 iModulon, we reapplied ICA on the Precise-1K dataset, now including the transcriptomics dataset from this study. This revealed new gene members to the iModulon containing the *rspAB* genes that we annotate as Rsp iModulon (Figure 4b). Notably, the Rsp iModulon includes the *betABIT* operon that encodes a choline transporter and oxidizing enzymes to take up and biosynthesize betaine from choline (Lamark et al., 1996). This is interesting because modulating osmolarity with betaine is a reported strategy for stabilizing recombinant proteins and improving production in *E. coli* (de Marco et al., 2005). Betaine is an effective osmolyte but can only be acquired from extracellular choline or taken up directly through the ProVWX transporter complex (Sévin and Sauer, 2014).

Both the Rsp and RpoH iModulons are activated in response to heterologous gene overexpression with a strong positive correlation (Figure 4c). To understand the connectivity between these responses, we explored whether other conditions from the Precise-1K library activate these iModulons. Across all conditions, those that show the highest activations of both iModulons are our overexpression conditions, particularly Cluster 1 samples (Red) and the BLG_T7 samples, and enzyme overexpression samples (orange) from Tan. et al. (Tan et al., 2020). However, the Rsp iModulon is not always activated with the RpoH iModulon. One enzyme, hsTPH_Ntermtrunc, shows activation of Rsp but not RpoH iModulon, suggesting that it may be folding properly but still stressing the Rsp iModulon response. Other stresses show activation of the RpoH iModulon without Rsp, specifically 44°C heat stress and 5% ethanol stress (Choudhary et al., 2020), suggesting that these are independent responses activated by different stimuli. Taken together, the Rsp iModulon is a characteristic that is mostly related to stress from overexpression of heterologous genes.

### 3.6 Alteration of the Rsp iModulon activity improves EGFP production

We further implemented different strategies to change Rsp iModulon activity to improve production of EGFP. To assess the effect of *rspAB* activation, we overexpressed *rspAB* genes using a constitutive promoter (Figure 5a). To evaluate the benefit of *betABIT* activation, we supplemented the cultures with either 5 mM choline or betaine at the time of rhamnose induction. The same cultivation conditions were used for generating the transcriptomics data, as described in Methods and Materials, and conditions were repeated for 4 replicates. By overexpressing *rspAB*, production of EGFP increased significantly (p<0.0005) by up to 32.2% with a slight reduction in growth rate (Figure 5a). By adding choline (5 mM), fluorescence levels increased significantly (p<0.05) by 9.5% (Figure 5b). Potentially, these strategies may be used to improve protein production for other proteins that activate the Rsp iModulon and may be applicable for achieving higher product titers in high-cell density settings.

**Figure 5.**
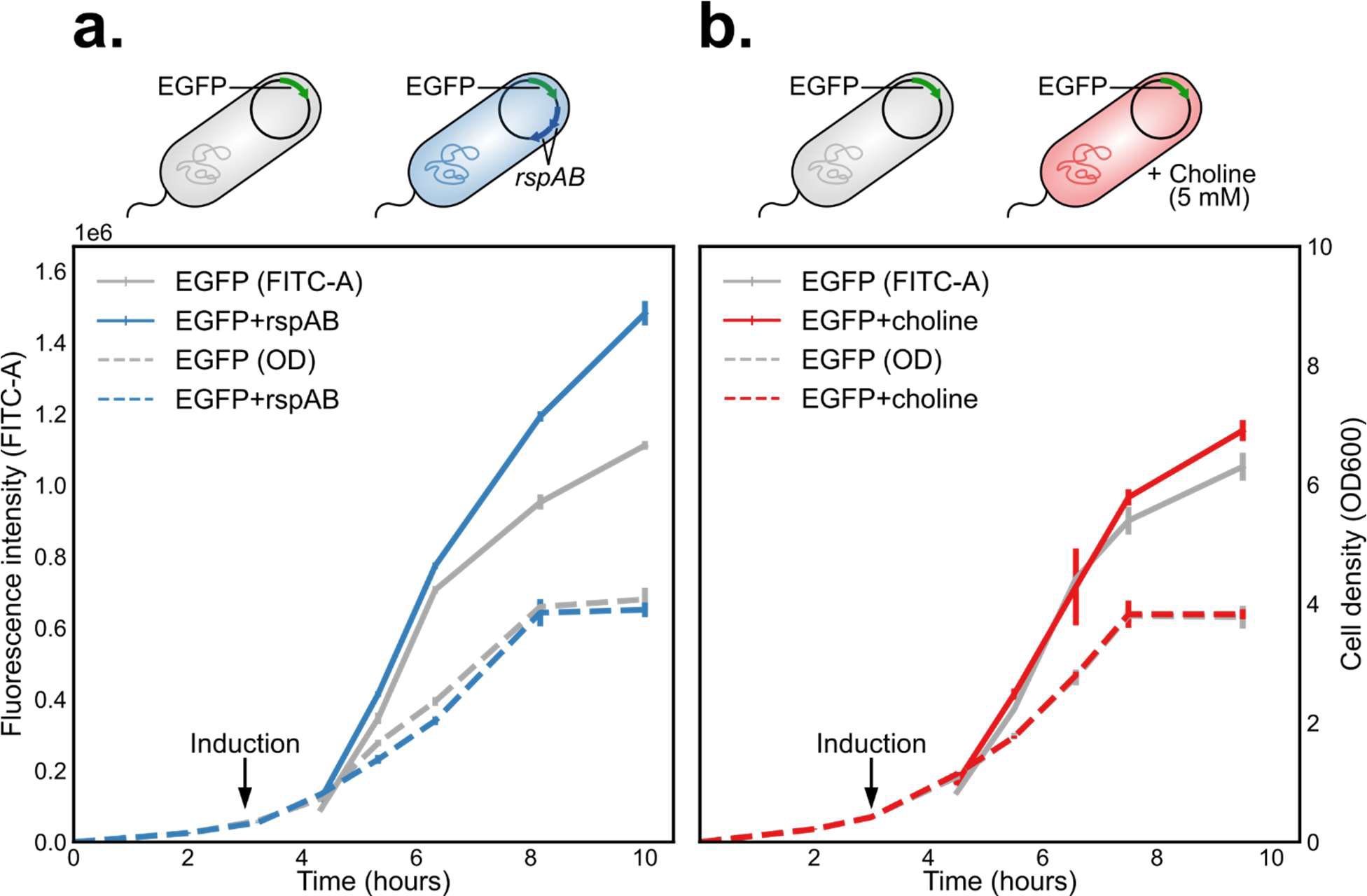
Changes to EGFP fluorescence (FITC-A) and growth (OD600) over time by **a.** *rspAB* co-overexpression and **b.** choline supplementation (5 mM). Cells were cultivated in shake flasks in 4 replicates and induced at OD = 0.40. Fluorescence levels were measured by flow cytometry.

The positive effect of *rspAB* expression on EGFP production may be attributed to downregulated RpoS regulon genes, which can be beneficial to protein production, as discussed in 3.2. Increased EGFP levels from co-overexpressing *rspAB* are most pronounced as cells enter the early stationary phase, which was also reported by Weikert et al. upon production of β-galactosidase (Weikert et al., 2000). One explanation that has been proposed is that RspA and RspB degrade homoserine lactone or homocysteine thiolactone that are signaling modulus activating RpoS (Goodrich-Blair and Kolter, 2000; Huisman and Kolter, 1994). These signal molecules accumulate after tRNAs become mis-activated with homoserine or homocysteine (Jakubowski, 1997, 1990), and this happens more frequently when concentrations of these amino acid intermediates are high (Jakubowski and Goldman, 1992). Possibly, translation of the heterologous gene may increase levels of amino acid intermediates, leading to increased misactivation levels which activate RpoS. We speculate that forcing the cells to stay in a ‘greed’ instead of ‘fear’ phenotype through RspAB may help allocate resources towards producing more protein products (Dalldorf et al., 2023). More recent studies indicate that RspA is a D-altronate dehydrogenase used in glucuronate metabolism (Wichelecki 2014, Gerlt 2005). However, there is no direct link between this part of metabolism and protein production. For future efforts, revealing the molecular function of RspAB may help identify other novel strategies for improving protein production.

Exogenously added choline, which is taken up and biosynthesized to betaine through the *betABIT* operon, is a novel strategy for improving protein production. Interestingly, direct betaine supplementation did not increase EGFP levels (Figure S7). One possible reason is that betaine is not taken up and accumulated by the cells. Betaine must be imported through another system encoded by the *proVWX* operon, which is captured by the Osmoprotectant iModulon. Overexpression samples in this study have slightly elevated activities of the Osmoprotectant iModulon, which is more highly activated by high NaCl conditions (Lennen et al., 2023). Studies that successfully improved protein production combined betaine supplementation with NaCl to stimulate betaine uptake (de Marco et al., 2005; Oganesyan et al., 2007). These results show that protein production can be improved through osmolyte supplementation but must be coordinated with the underlying regulatory network to have a positive effect.

## 4. Conclusion

Stress responses to the overexpression of 10 different heterologous genes were studied at the transcriptional level in an *E. coli* production host. These responses were found to depend on the characteristics of the heterologous gene. Transcriptomes taken during overexpression of different heterologous genes were categorized into two different clusters, separating them from the control samples. To interpret the transcriptional response in these two clusters, iModulons activities in our transcriptomics dataset was computed by applying the Precise-1K RNAseq knowledge base. This revealed that one cluster of overexpression samples activates a response characterized by the heat shock stress iModulon, RpoH, and an uncharacterized iModulon UC-9. The other cluster of overexpression samples activate more diverse responses, but all share the activation of the Cold Shock and gcvB sRNA iModulons. To further understand and identify other genes related to the uncharacterized UC-9 response, we included our dataset for ICA. We then designed genetic modifications and media supplementations to complement the genes in this iModulon, which were shown experimentally to improve the production of EGFP. Our study has demonstrated that ICA and iModulon analysis of transcriptomics data is a promising use of big data analytics to identify novel traits of host stress responses that can guide the design for better protein production.

## Software

Illustrations used in Figure 1, Figure 5, Figure S4, and Figure S7 were created with BioRender.com.

## Author Contributions

C.R., E.Ö., L.Y., and B.O.P. conceived the study. C.R., E.Ö., L.Y., B.O.P., L.J.J., and M.N. designed the experiments. C.R. generated the data. C.R. E.Ö. and L.Y. analyzed the data. C.R, F.B., and M.J. performed the computational preprocessing. C.R., B.O.P., E.Ö., and L.Y. wrote the manuscript with contributions from all the other co-authors.

## Acknowledgments

This work was funded by the Novo Nordisk Foundation (Grant Number NNF20CC0035580). We thank Se Hyeuk Kim, Peter Ehlert Jensen, Andreas Birk Bertelsen, Cristina Hernández Rollán, Anargyros Alexiou, Christina Lenhard, Lena Elisabeth Charlotte Heer, and Arsenios Vlassis for assistance in the experiments, and Daniel Zielinski for valuable input on the manuscript.

## Supplemental material

### 6. Supplemental Tables and Figures

**Table S1.** Composition of trace metal solution used in this study.

| Trace metal solution recipe |  |
| --- | --- |
| Component | Concentration (g/L) |
| CuSO<sub>4</sub>·5H<sub>2</sub>O | 0.48 |
| CoCl<sub>2</sub>·6H<sub>2</sub>O | 0.54 |
| ZnSO<sub>4</sub>·7H<sub>2</sub>O | 0.54 |
| CaCl<sub>2</sub>·2H<sub>2</sub>O | 2 |
| FeCl<sub>3</sub>·6H<sub>2</sub>O | 41.8 |
| MnSO<sub>4</sub>·H<sub>2</sub>O | 0.3 |
| Na<sub>2</sub> EDTA·2H<sub>2</sub>O | 33.4 |

**Figure S1.**
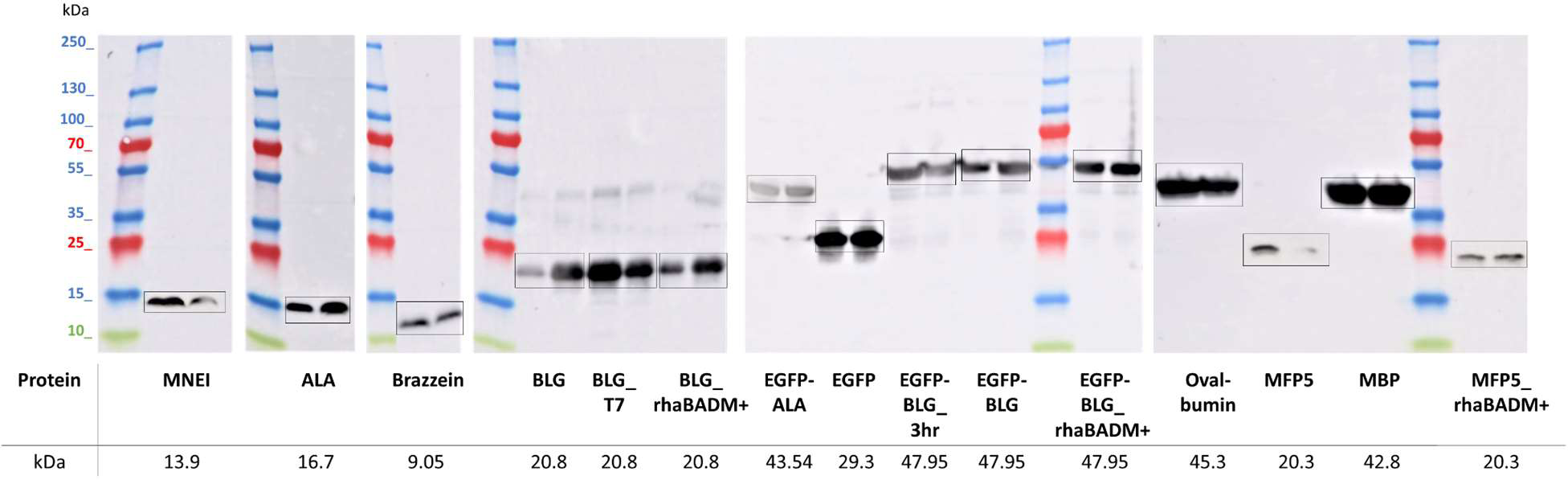
Western blots in heterologous overexpression samples. All proteins were successfully produced. A larger band (∼40kDA) was identified during the expression of BLG, which is likely BLG homodimers.

**Figure S2.**
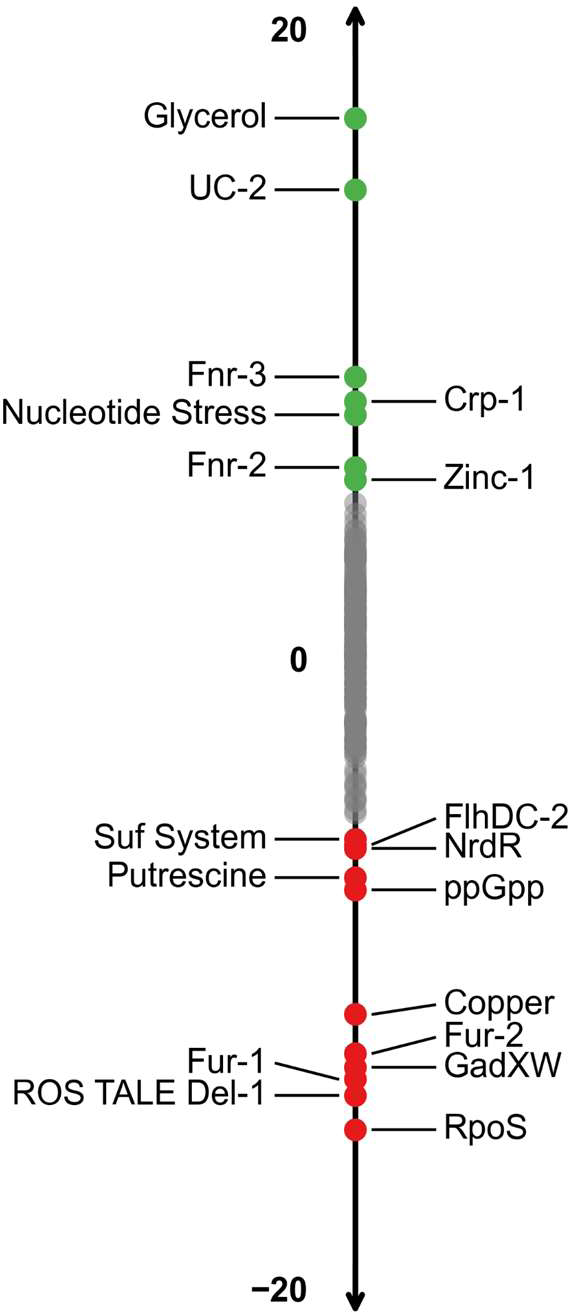
iModulon activities in control samples from this study benchmarked with Precise-1K. The mean activities of iModulons in control samples (cluster 3, see Figure 2a) are compared to wildtype control samples in the Precise-1K RNA-seq compendium (MG1655 grown in M9 glucose, control wt_glc 1 and control wt_glc 2). Wildtype controls are used as the baseline for iModulon calculations, and their gene expression levels are set to 0 by default. Green and red iModulons are significantly elevated or lowered, respectively with a iModulon activity threshold of 5 and a false detection rate set to 0.1. Many differentially activated iModulons are related to medium composition. Glycerol iModulon is significantly upregulated since glycerol is used as a carbon source. Iron regulation iModulons Fur-1 and Fur-2 are significantly downregulated, indicating sufficiently high iron levels in the medium used. Similarly, Copper iModulon is downregulated, but Zinc-1 is upregulated, indicating slight starvation of Zinc. Cells are more anaerobically stressed, as shown by increased levels of Fnr-2 and Fnr-3 iModulons. The stringent response is inactive, shown by lowered levels of RpoS and ppGpp iModulons.

**Figure S3.**
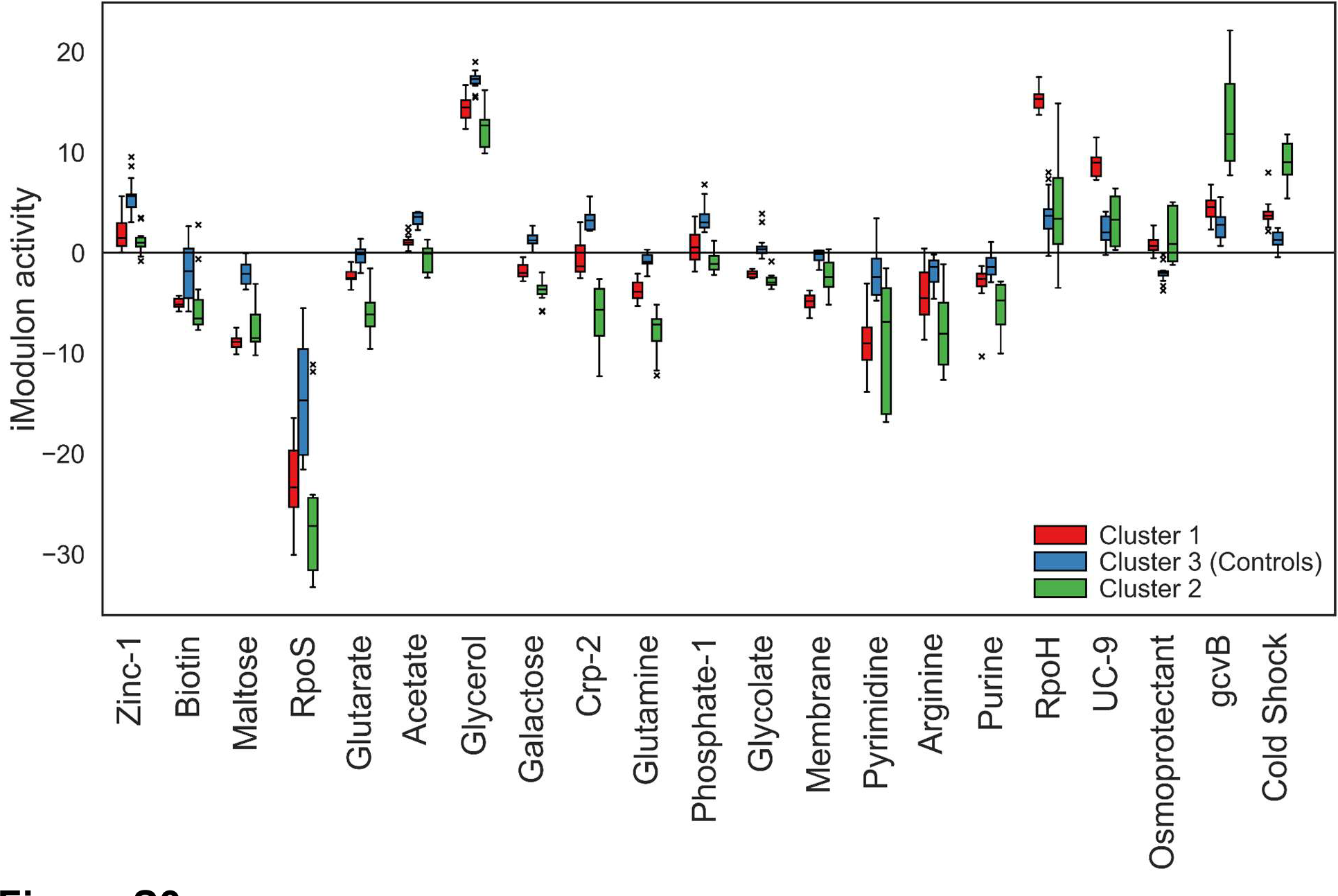
Boxplot of 21 most differentially activated iModulons during gene overexpression (Clusters 1 and 2, see Figure 2a), compared to controls (Cluster 3). RpoH and UC-9 iModulons are activated in Cluster 1 overexpression samples, compared to Cluster 3 control samples. gcvB and Cold Shock iModulons are activated in Cluster 2.

**Figure S4.**
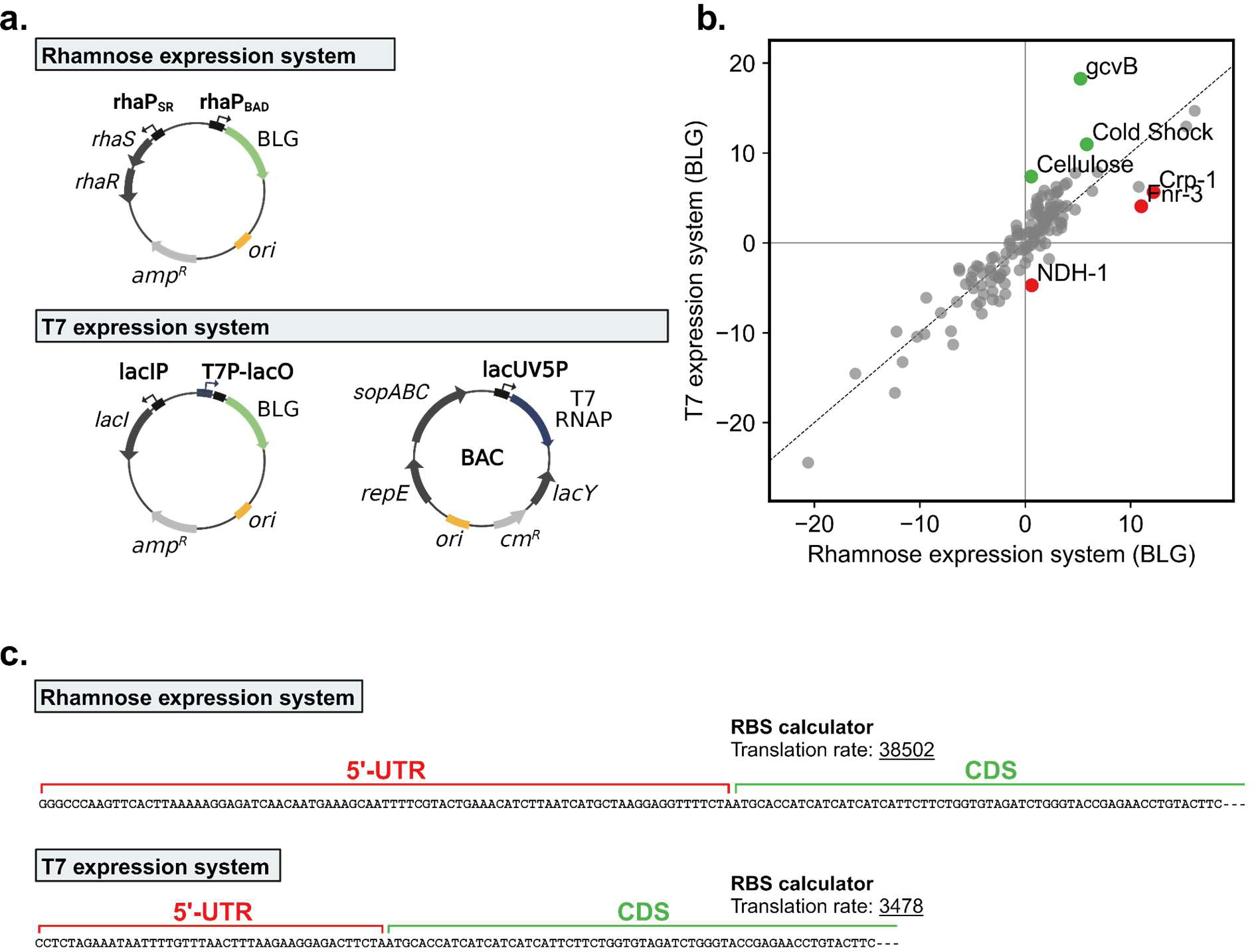
Comparison of rhamnose and T7 RNA Polymerase (T7 RNAP) expression systems for BLG overexpression. **a.** Overview of plasmids used for the rhamnose system and for the T7 RNAP expression system. The T7 RNAP system yields higher levels of heterologous mRNA. **b.** Differential iModulon Activity plot of BLG expressed using the T7 RNAP system against BLG using the rhamnose system. Significance is determined using standard threshold (5) and false detection rate (0.1) settings. The gcvB, Cold Shock, and Cellulose iModulons are upregulated during strong expression using T7 RNAP, while energy metabolism (NDH-1 iModulon) and cAMP receptor protein regulation (Crp-1 iModulon) are downregulated. **c.** Calculation of translation initiation rates using the Salis Lab RBS calculator (v.2.1) from Salis Lab (Reis and Salis, 2020). The rhamnose expression system was estimated to have a 30 times higher translation rate than the T7-based expression system.

**Figure S5.**
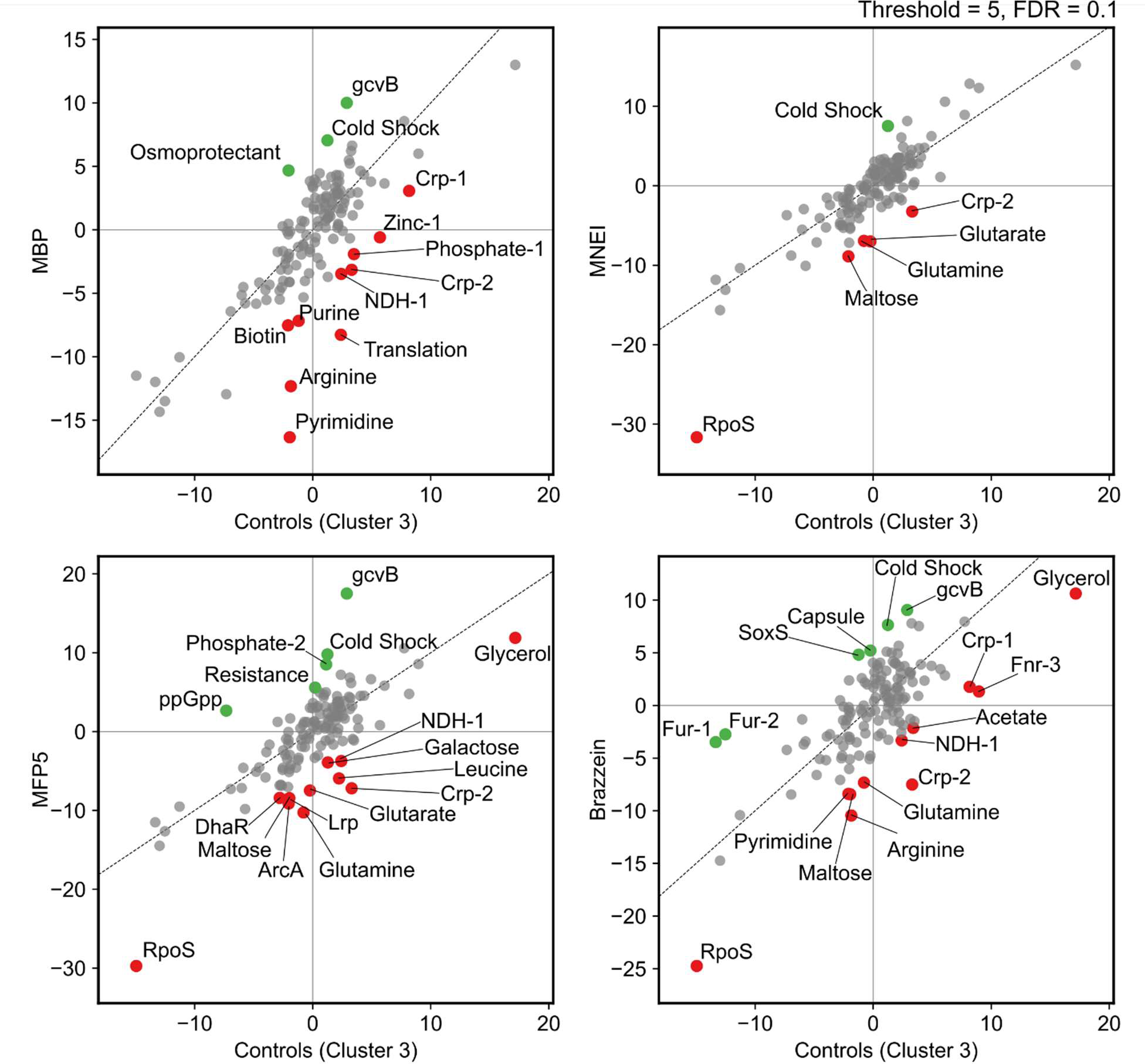
Differential iModulon activity during overexpression of MBP, MFP5, MNEI, and Brazzein showing diverse transcriptional responses. Results are presented as Differential iModulon Activity plots, comparing iModulon activities with control conditions (Cluster 3, see Figure 2a). MBP, encoded by *malE* in *E. coli*, is a protein with excellent folding behavior and hence is used as a solubility tag for protein production. Overexpression of MBP activated the Osmoprotectant iModulon, a small iModulon comprised of the *proVWX* operon, encoding a transporter for betaine and proline (Lamark et al., 1996). Regulation of other functions was greatly lowered during the expression of MBP, particularly ribosome biogenesis (Translation iModulon), nucleotide metabolism (Pyrimidine and Purine iModulons), and arginine metabolism. MFP5 is an artificial dimer that is highly rich in tyrosine (23%), glycine (19%) and lysine (18%). Overexpression of the MFP5 gene reduced growth and yielded the lowest observed biomass of any of the tested heterologous genes (see Figure 1). The transcriptional response is characterized by upregulated ribosome biogenesis (Translation and ppGpp iModulons) and phosphate metabolism (Phosphate-2 iModulon). Several amino acid iModulons were downregulated (Lrp, Leucine, and Glutamine iModulons). MNEI is a small protein described to have strong folding stability (Leone et al., 2016). Overexpression of the MNEI gene also resulted in lowered growth, however, iModulon activities changed little compared to control conditions. Brazzein is a cysteine-rich protein that triggers an oxidative stress-related stress response reflected by activation of the SoxS iModulon and iron uptake (Fur-1 and Fur-2 iModulons).

**Figure S6.**
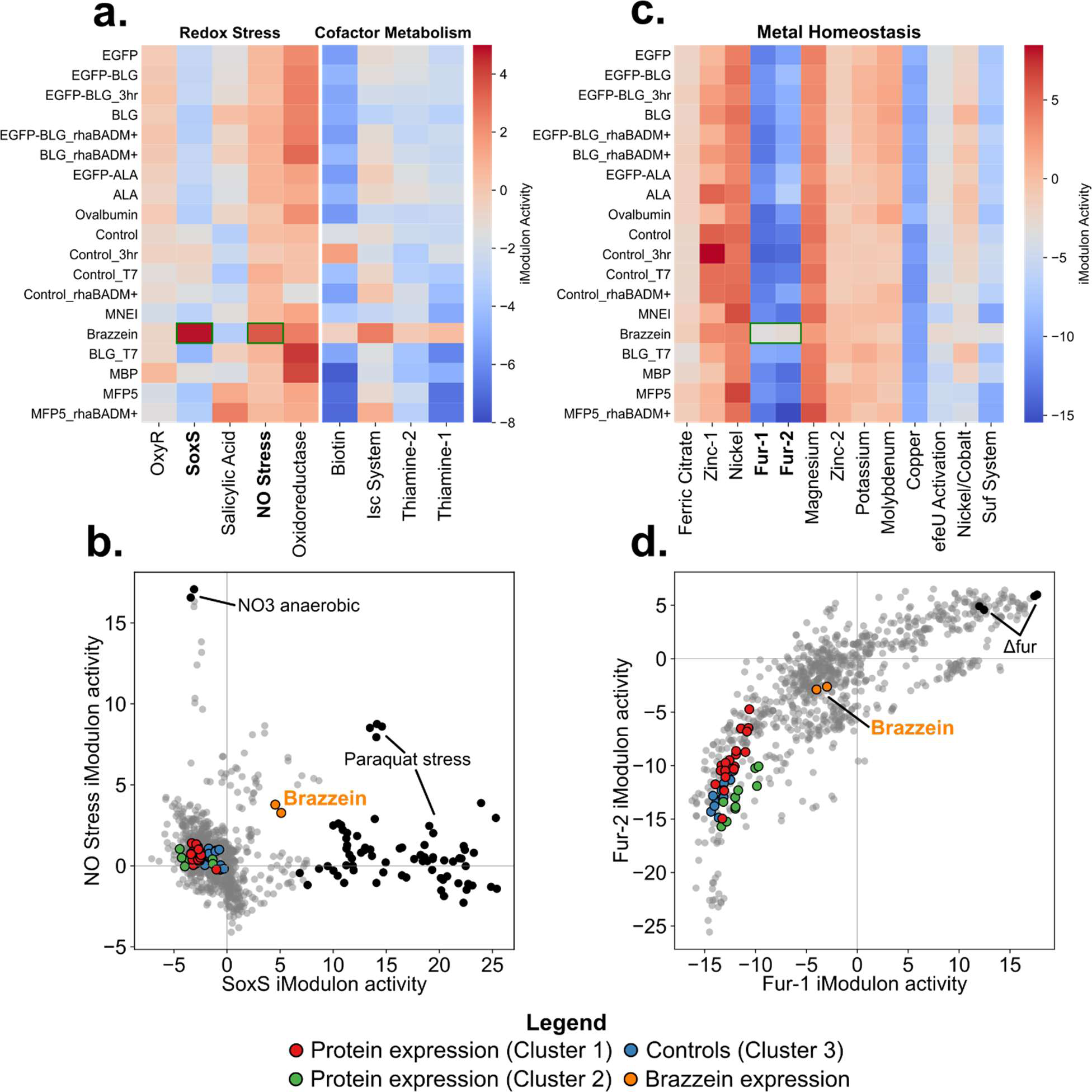
Brazzein overexpression activates a redox and iron uptake stress response. **a.** Two iModulons related to reactive oxygen species detoxification are particularly high in the brazzein expression samples; the SoxS and NO Stress (Nitric Oxide) iModulons (green boxes). **b.** The activities of these two iModulons with all other samples in the Precise-1K RNAseq library in a phase plane. SoxS responds to superoxide stress (O2^-^) and can otherwise be activated by adding paraquat, a toxic oxidant. The NO Stress iModulon responds to nitric oxide radicals (^•^NO) and activates during anaerobic growth on nitrate. Interestingly, the OxyR iModulon is not activated, which is an important oxidative stress response to hydrogen peroxide (H2O2). **c.** The redox stress during Brazzein overexpression is accompanied by activation of iron uptake iModulons Fur-1 and Fur-2. These iModulons encode genes related to siderophore synthesis, iron transport, and hydrolysis systems. **d.** The activation of Fur-1 and Fur-2 iModulons follows the non-linear correlation that has been described earlier(Rychel et al., 2022; Sastry et al., 2021). Iron-sulfur (Fe-S) cluster biogenesis was also found upregulated which includes the housekeeping Isc System (a.), and the Suf pathway (c.), which is active under oxidative stress (Ayala-Castro et al., 2008). We speculate that the accumulation of cysteine-rich Brazzein makes the host more sensitive to oxidative stress. Cysteines are the first residues on proteins to become oxidized, which could desensitize the finely tuned levels of redox-sensitive proteins, such as OxyR (Leichert et al., 2008; Lushchak, 2001). The increased concentration of cysteines may interact with intracellular iron ions, triggering iron uptake.

**Figure S7.**
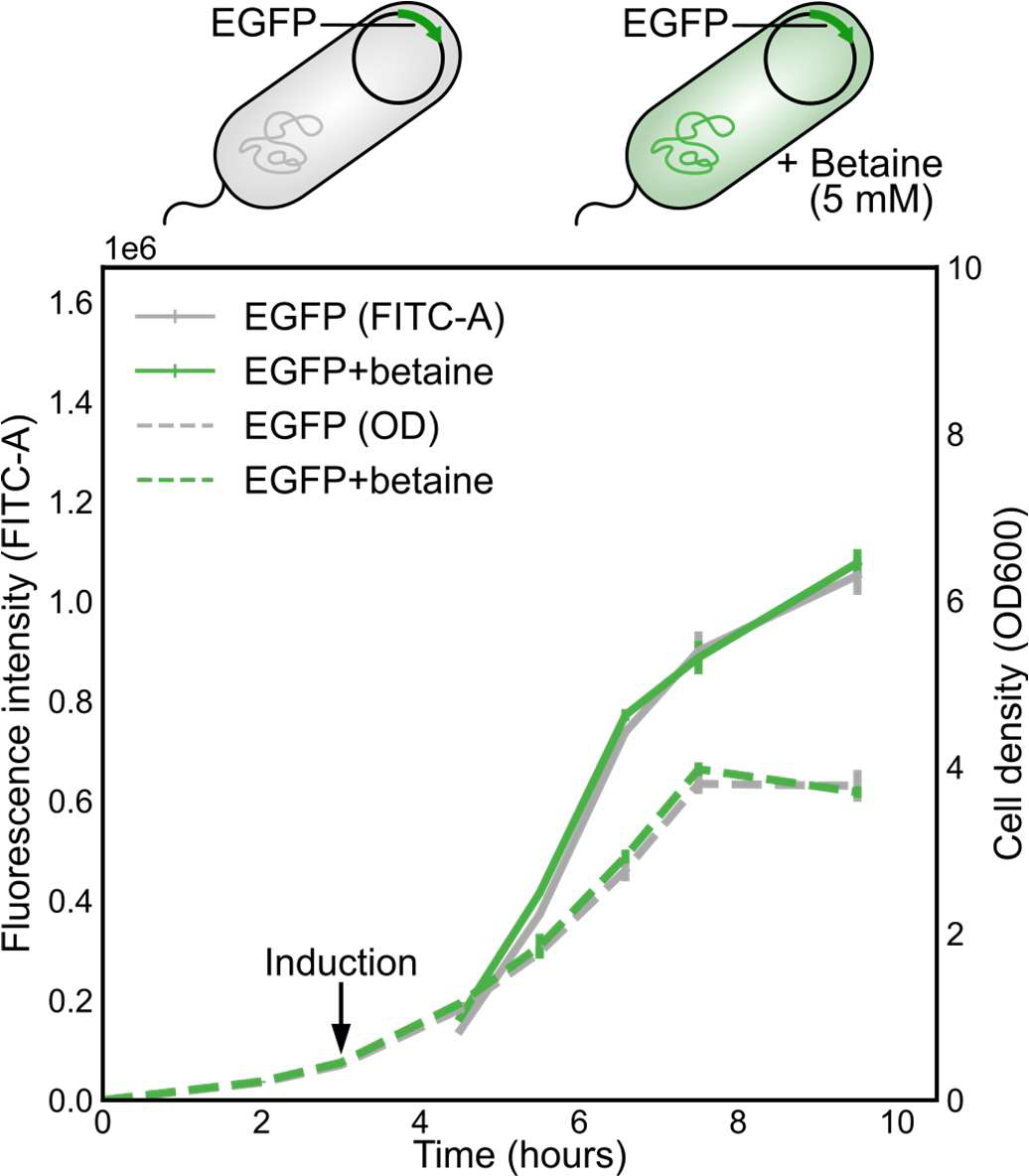
EGFP fluorescence (FITC-A) and growth (OD600) over time during betaine supplementation (5 mM). Cells were grown in shake flasks for 4 biological replicates, and fluorescence levels were taken by flow cytometry. Betaine supplementation had no significant effect on fluorescence intensity over time or growth.

## Notes

### Competing Interest Statement

The authors have declared no competing interest.

## Literature

1. Arnau, J., Yaver, D., Hjort, C.M., 2020. Strategies and Challenges for the Development of Industrial Enzymes Using Fungal Cell Factories, in: Nevalainen, H. (Ed.), Grand Challenges in Fungal Biotechnology. Springer International Publishing, Cham, pp. 179–210. 10.1007/978-3-030-29541-7_7

2. Arsène, F., Tomoyasu, T., Bukau, B., 2000. The heat shock response of Escherichia coli. Int. J. Food Microbiol. 55, 3–9. 10.1016/s0168-1605(00)00206-3

3. Augustin, M.A., Hartley, C.J., Maloney, G., Tyndall, S., 2023. Innovation in precision fermentation for food ingredients. Crit. Rev. Food Sci. Nutr. 1–21. 10.1080/10408398.2023.2166014

4. Ayala-Castro, C., Saini, A., Outten, F.W., 2008. Fe-S cluster assembly pathways in bacteria. Microbiol. Mol. Biol. Rev. 72, 110–25, table of contents. 10.1128/MMBR.00034-07

5. Behdenna, A., Colange, M., Haziza, J., Gema, A., Appé, G., Azencott, C.-A., Nordor, A., 2023. pyComBat, a Python tool for batch effects correction in high-throughput molecular data using empirical Bayes methods. bioRxiv. 10.1101/2020.03.17.995431

6. Bhatwa, A., Wang, W., Hassan, Y.I., Abraham, N., Li, X.-Z., Zhou, T., 2021. Challenges Associated With the Formation of Recombinant Protein Inclusion Bodies in Escherichia coli and Strategies to Address Them for Industrial Applications. Front Bioeng Biotechnol 9, 630551. 10.3389/fbioe.2021.630551

7. Chen, H., Previero, A., Deutscher, M.P., 2019. A novel mechanism of ribonuclease regulation: GcvB and Hfq stabilize the mRNA that encodes RNase BN/Z during exponential phase. J. Biol. Chem. 294, 19997–20008. 10.1074/jbc.RA119.011367

8. Chen, Y., Banerjee, D., Mukhopadhyay, A., Petzold, C.J., 2020. Systems and synthetic biology tools for advanced bioproduction hosts. Curr. Opin. Biotechnol. 64, 101–109. 10.1016/j.copbio.2019.12.007

9. Choudhary, K.S., Kleinmanns, J.A., Decker, K., Sastry, A.V., Gao, Y., Szubin, R., Seif, Y., Palsson, B.O., 2020. Elucidation of Regulatory Modes for Five Two-Component Systems in Escherichia coli Reveals Novel Relationships. mSystems 5. 10.1128/mSystems.00980-20

10. Dalldorf, C., Rychel, K., Szubin, R., Hefner, Y., Patel, A., Zielinski, D., Palsson, B., 2023. The hallmarks of a tradeoff in transcriptomes that balances stress and growth functions. Res Sq. 10.21203/rs.3.rs-2729651/v1

11. Danner, E.W., Kan, Y., Hammer, M.U., Israelachvili, J.N., Waite, J.H., 2012. Adhesion of Mussel Foot Protein Mefp-5 to Mica: An Underwater Superglue. Biochemistry 51, 6511–6518. 10.1021/bi3002538

12. Davy, A.M., Kildegaard, H.F., Andersen, M.R., 2017. Cell Factory Engineering. Cell Syst 4, 262–275. 10.1016/j.cels.2017.02.010

13. de Marco, A., Vigh, L., Diamant, S., Goloubinoff, P., 2005. Native folding of aggregation-prone recombinant proteins in Escherichia coli by osmolytes, plasmid- or benzyl alcohol-overexpressed molecular chaperones. Cell Stress Chaperones 10, 329–339. 10.1379/csc-139r.1

14. Dürrschmid, K., Reischer, H., Schmidt-Heck, W., Hrebicek, T., Guthke, R., Rizzi, A., Bayer, K., 2008. Monitoring of transcriptome and proteome profiles to investigate the cellular response of E. coli towards recombinant protein expression under defined chemostat conditions. J. Biotechnol. 135, 34–44. 10.1016/j.jbiotec.2008.02.013

15. Eskandari, A., Leow, T.C., Rahman, M.B.A., Oslan, S.N., 2020. Antifreeze Proteins and Their Practical Utilization in Industry, Medicine, and Agriculture. Biomolecules 10. 10.3390/biom10121649

16. Giuliodori, A.M., 2016. Cold shock response in Escherichia coli: A model system to study posttranscriptional regulation, in: Stress and Environmental Regulation of Gene Expression and Adaptation in Bacteria. John Wiley & Sons, Inc., Hoboken, NJ, USA, pp. 859–872. 10.1002/9781119004813.ch84

17. Goodrich-Blair, H., Kolter, R., 2000. Homocysteine thiolactone is a positive effector of σS levels in Escherichia coli. FEMS Microbiol. Lett. 185, 117–121. 10.1111/j.1574-6968.2000.tb09048.x

18. Haddadin, F.T., Harcum, S.W., 2005. Transcriptome profiles for high-cell-density recombinant and wild-type Escherichia coli. Biotechnol. Bioeng. 90, 127–153. 10.1002/bit.20340

19. Huisman, G.W., Kolter, R., 1994. Sensing starvation: a homoserine lactone--dependent signaling pathway in Escherichia coli. Science 265, 537–539. 10.1126/science.7545940

20. Jakubowski, H., 1997. Aminoacyl thioester chemistry of class II aminoacyl-tRNA synthetases. Biochemistry 36, 11077–11085. 10.1021/bi970589n

21. Jakubowski, H., 1990. Proofreading in vivo: editing of homocysteine by methionyl-tRNA synthetase in Escherichia coli. Proc. Natl. Acad. Sci. U. S. A. 87, 4504–4508. 10.1073/pnas.87.12.4504

22. Jakubowski, H., Goldman, E., 1992. Editing of errors in selection of amino acids for protein synthesis. Microbiol. Rev. 56, 412–429. 10.1128/mr.56.3.412-429.1992

23. Jeong, K.J., Choi, J.H., Yoo, W.M., Keum, K.C., Yoo, N.C., Lee, S.Y., Sung, M.-H., 2004. Constitutive production of human leptin by fed-batch culture of recombinant rpoS-Escherichia coli. Protein Expr. Purif. 36, 150–156. 10.1016/j.pep.2004.04.007

24. Jiang, W., Jones, P., Inouye, M., 1993. Chloramphenicol induces the transcription of the major cold shock gene of Escherichia coli, cspA. J. Bacteriol. 175, 5824–5828. 10.1128/jb.175.18.5824-5828.1993

25. Jones, P.G., Inouye, M., 1994. The cold-shock response--a hot topic. Mol. Microbiol. 11, 811–818. 10.1111/j.1365-2958.1994.tb00359.x

26. Kim, E., Dai, B., Qiao, J.B., Li, W., Fortner, J.D., Zhang, F., 2018. Microbially Synthesized Repeats of Mussel Foot Protein Display Enhanced Underwater Adhesion. ACS Appl. Mater. Interfaces 10, 43003–43012. 10.1021/acsami.8b14890

27. Kirk, O., Borchert, T.V., Fuglsang, C.C., 2002. Industrial enzyme applications. Curr. Opin. Biotechnol. 13, 345–351. 10.1016/s0958-1669(02)00328-2

28. Lalaouna, D., Eyraud, A., Devinck, A., Prévost, K., Massé, E., 2019. GcvB small RNA uses two distinct seed regions to regulate an extensive targetome. Mol. Microbiol. 111, 473–486. 10.1111/mmi.14168

29. Lamark, T., Røkenes, T.P., McDougall, J., Strøm, A.R., 1996. The complex bet promoters of Escherichia coli: regulation by oxygen (ArcA), choline (BetI), and osmotic stress. J. Bacteriol. 178, 1655–1662. 10.1128/jb.178.6.1655-1662.1996

30. Lamoureux, C.R., Decker, K.T., Sastry, A.V., Rychel, K., Gao, Y., McConn, J.L., Zielinski, D.C., Palsson, B.O., 2023. A multi-scale expression and regulation knowledge base for Escherichia coli. Nucleic Acids Res. 10.1093/nar/gkad750

31. Leichert, L.I., Gehrke, F., Gudiseva, H.V., Blackwell, T., Ilbert, M., Walker, A.K., Strahler, J.R., Andrews, P.C., Jakob, U., 2008. Quantifying changes in the thiol redox proteome upon oxidative stress in vivo. Proc. Natl. Acad. Sci. U. S. A. 105, 8197–8202. 10.1073/pnas.0707723105

32. Lennen, R.M., Lim, H.G., Jensen, K., Mohammed, E.T., Phaneuf, P.V., Noh, M.H., Malla, S., Börner, R.A., Chekina, K., Özdemir, E., Bonde, I., Koza, A., Maury, J., Pedersen, L.E., Schöning, L.Y., Sonnenschein, N., Palsson, B.O., Nielsen, A.T., Sommer, M.O.A., Herrgård, M.J., Feist, A.M., 2023. Laboratory evolution reveals general and specific tolerance mechanisms for commodity chemicals. Metab. Eng. 76, 179–192. 10.1016/j.ymben.2023.01.012

33. Leone, S., Pica, A., Merlino, A., Sannino, F., Temussi, P.A., Picone, D., 2016. Sweeter and stronger: enhancing sweetness and stability of the single chain monellin MNEI through molecular design. Sci. Rep. 6, 1–10. 10.1038/srep34045

34. Lieder, S., Nikel, P.I., de Lorenzo, V., Takors, R., 2015. Genome reduction boosts heterologous gene expression in Pseudomonas putida. Microb. Cell Fact. 14, 23. 10.1186/s12934-015-0207-7

35. Lushchak, V.I., 2001. Oxidative Stress and Mechanisms of Protection Against It in Bacteria. Biochemistry 66, 476–489. 10.1023/A:1010294415625

36. McConn, J.L., Lamoureux, C.R., Poudel, S., Palsson, B.O., Sastry, A.V., 2021. Optimal dimensionality selection for independent component analysis of transcriptomic data. BMC Bioinformatics 22, 584. 10.1186/s12859-021-04497-7

37. Ming, D., Hellekant, G., 1994. Brazzein, a new high-potency thermostable sweet protein from Pentadiplandra brazzeana B. FEBS Lett. 355, 106–108. 10.1016/0014-5793(94)01184-2

38. Miserez, A., Yu, J., Mohammadi, P., 2023. Protein-Based Biological Materials: Molecular Design and Artificial Production. Chem. Rev. 123, 2049–2111. 10.1021/acs.chemrev.2c00621

39. Oganesyan, N., Ankoudinova, I., Kim, S.-H., Kim, R., 2007. Effect of osmotic stress and heat shock in recombinant protein overexpression and crystallization. Protein Expr. Purif. 52, 280–285. 10.1016/j.pep.2006.09.015

40. Oh, M.K., Liao, J.C., 2000. DNA microarray detection of metabolic responses to protein overproduction in Escherichia coli. Metab. Eng. 2, 201–209. 10.1006/mben.2000.0149

41. Ow, D.S.-W., Lim, D.Y.-X., Nissom, P.M., Camattari, A., Wong, V.V.-T., 2010. Co-expression of Skp and FkpA chaperones improves cell viability and alters the global expression of stress response genes during scFvD1.3 production. Microb. Cell Fact. 9, 22. 10.1186/1475-2859-9-22

42. Phadtare, S., 2004. Recent developments in bacterial cold-shock response. Curr. Issues Mol. Biol. 6, 125–136.

43. Pouresmaeil, M., Azizi-Dargahlou, S., 2023. Factors involved in heterologous expression of proteins in E. coli host. Arch. Microbiol. 205, 212. 10.1007/s00203-023-03541-9

44. Puetz, J., Wurm, F.M., 2019. Recombinant Proteins for Industrial versus Pharmaceutical Purposes: A Review of Process and Pricing. Processes 7, 476. 10.3390/pr7080476

45. Reis, A.C., Salis, H.M., 2020. An Automated Model Test System for Systematic Development and Improvement of Gene Expression Models. ACS Synth. Biol. 9, 3145–3156. 10.1021/acssynbio.0c00394

46. Rettenbacher, L.A., Arauzo-Aguilera, K., Buscajoni, L., Castillo-Corujo, A., Ferrero-Bordera, B., Kostopoulou, A., Moran-Torres, R., Núñez-Nepomuceno, D., Öktem, A., Palma, A., Pisent, B., Puricelli, M., Schilling, T., Tungekar, A.A., Walgraeve, J., Humphreys, D., von der Haar, T., Gasser, B., Mattanovich, D., Ruddock, L., van Dijl, J.M., 2022. Microbial protein cell factories fight back? Trends Biotechnol. 40, 576–590. 10.1016/j.tibtech.2021.10.003

47. Rong, Y., Jensen, S.I., Lindorff-Larsen, K., Nielsen, A.T., 2023. Folding of heterologous proteins in bacterial cell factories: Cellular mechanisms and engineering strategies. Biotechnol. Adv. 63, 108079. 10.1016/j.biotechadv.2022.108079

48. Rychel, K., Decker, K., Sastry, A.V., Phaneuf, P.V., Poudel, S., Palsson, B.O., 2021. iModulonDB: a knowledgebase of microbial transcriptional regulation derived from machine learning. Nucleic Acids Res. 49, D112–D120. 10.1093/nar/gkaa810

49. Rychel, K., Tan, J., Patel, A., Lamoureux, C., Hefner, Y., Szubin, R., Johnsen, J., Mohamed, E.T.T., Phaneuf, P.V., Anand, A., Olson, C.A., Park, J.H., Sastry, A.V., Yang, L., Feist, A.M., Palsson, B.O., 2022. Lab evolution, transcriptomics, and modeling reveal mechanisms of paraquat tolerance. bioRxiv. 10.1101/2022.12.20.521246

50. Sastry, A.V., Dillon, N., Anand, A., Poudel, S., Hefner, Y., Xu, S., Szubin, R., Feist, A.M., Nizet, V., Palsson, B., 2021. Machine Learning of Bacterial Transcriptomes Reveals Responses Underlying Differential Antibiotic Susceptibility. mSphere 6, e0044321. 10.1128/mSphere.00443-21

51. Sastry, A.V., Gao, Y., Szubin, R., Hefner, Y., Xu, S., Kim, D., Choudhary, K.S., Yang, L., King, Z.A., Palsson, B.O., 2019. The Escherichia coli transcriptome mostly consists of independently regulated modules. Nat. Commun. 10, 1–14. 10.1038/s41467-019-13483-w

52. Sévin, D.C., Sauer, U., 2014. Ubiquinone accumulation improves osmotic-stress tolerance in Escherichia coli. Nat. Chem. Biol. 10, 266–272. 10.1038/nchembio.1437

53. Sharma, A.K., Mahalik, S., Ghosh, C., Singh, A.B., Mukherjee, K.J., 2011. Comparative transcriptomic profile analysis of fed-batch cultures expressing different recombinant proteins in Escherichia coli. AMB Express 1, 33. 10.1186/2191-0855-1-33

54. Shen, S., Packer, H., 2022. Codon optimization tool makes synthetic gene design easy [WWW Document]. Integrated DNA Technologies. URL https://eu.idtdna.com/pages/education/decoded/article/idt-codon-optimization-tool-makes-synthetic-gene-design-easy (accessed 8.29.23).

55. Tan, J., Sastry, A.V., Fremming, K.S., Bjørn, S.P., Hoffmeyer, A., Seo, S., Voldborg, B.G., Palsson, B.O., 2020. Independent component analysis of E. coli’s transcriptome reveals the cellular processes that respond to heterologous gene expression. Metab. Eng. 61, 360–368. 10.1016/j.ymben.2020.07.002

56. Tyo, K.E.J., Liu, Z., Petranovic, D., Nielsen, J., 2012. Imbalance of heterologous protein folding and disulfide bond formation rates yields runaway oxidative stress. BMC Biol. 10, 16. 10.1186/1741-7007-10-16

57. Weikert, C., Canonaco, F., Sauer, U., Bailey, J.E., 2000. Co-overexpression of RspAB improves recombinant protein production in Escherichia coli. Metab. Eng. 2, 293–299. 10.1006/mben.2000.0163

58. Wichelecki, D.J., Balthazor, B.M., Chau, A.C., Vetting, M.W., Fedorov, A.A., Fedorov, E.V., Lukk, T., Patskovsky, Y.V., Stead, M.B., Hillerich, B.S., Seidel, R.D., Almo, S.C., Gerlt, J.A., 2014. Discovery of function in the enolase superfamily: D-mannonate and d-gluconate dehydratases in the D-mannonate dehydratase subgroup. Biochemistry 53, 2722–2731. 10.1021/bi500264p

59. Ziegler, M., Zieringer, J., Döring, C.-L., Paul, L., Schaal, C., Takors, R., 2021. Engineering of a robust Escherichia coli chassis and exploitation for large-scale production processes. Metab. Eng. 67, 75–87. 10.1016/j.ymben.2021.05.011

